# Integrative computational toxicology reveals PFOS and PFHxS associated inflammatory keratinocyte niches in psoriasis through exposure transcriptomics, single-cell spatial mapping and token-aware virtual perturbation

**DOI:** 10.64898/2026.07.09.737426

**Authors:** Junhao Ma, Junjie Tan, Yucan Wang, Shihao Zhou, Shanshan Cai, Qianying Yu

## Abstract

Per- and polyfluoroalkyl substances (PFAS) are persistent toxicants with immunological, metabolic and epithelial effects, but their relevance to inflammatory skin disease remains unclear. We developed a computational toxicology framework to test whether perfluoroalkyl sulfonate programs, especially perfluorooctanesulfonic acid (PFOS) and perfluorohexanesulfonic acid (PFHxS), converge with psoriasis-associated keratinocyte inflammation. Exposure transcriptomes were derived from GSE236956, in which human embryonic stem cell-derived epithelial-lineage models were exposed to 10 μM PFAS for 8–16 days. Six PFAS were prioritized using descriptors, Tanimoto similarity, toxicology evidence, adverse outcome pathway (AOP)-like key events, exposure differentially expressed gene burden and read-across support. PFAS signatures were integrated with psoriasis bulk transcriptomes, single-cell RNA sequencing, keratinocyte-state mapping, regulator and communication inference, spatial transcriptomics and token-aware Geneformer-compatible virtual perturbation. PFOS ranked highest in integrated prioritization, followed by PFHxS and perfluorooctanoic acid. PFHxS produced a smaller but directionally informative signature within a PFOS-dominant perfluoroalkyl sulfonate footprint. The shared PFOS and PFHxS program converged with psoriasis through inflammatory keratinocyte, epidermal-stress, cytoskeletal and lipid-related modules. Single-cell and spatial analyses localized the program to activated keratinocytes and inflammatory epidermal niches, with strong spatial co-localization with inflammatory keratinocyte and epidermal stress scores. Virtual perturbation prioritized S100A9, S100A8, KRT16, IL36G, CCL20, CXCL8, FABP5, KRT17, FOS, JUN and NFKBIZ as candidate effectors. These findings support an exposure-informed, experimentally testable hypothesis linking persistent perfluoroalkyl sulfonate programs to keratinocyte inflammatory niches in psoriasis.

## 1. Introduction

Psoriasis is a chronic immune-mediated inflammatory skin disease characterized by epidermal hyperplasia, abnormal keratinocyte differentiation, vascular remodeling and immune-cell infiltration. Although the interleukin (IL)-23/IL-17 axis remains central to current mechanistic and therapeutic models, psoriasis cannot be explained by cytokine signaling alone. Keratinocytes actively shape lesional inflammation by producing chemokines, antimicrobial peptides, alarmins, lipid mediators and stress-response molecules that recruit and activate immune and stromal compartments (Nestle et al., 2009; Rendon and Schäkel, 2019). Clinical reviews also describe psoriasis as a coordinated epithelial-immune disease involving epidermal activation, immune recruitment and vascular remodeling (Boehncke and Schön, 2015; Armstrong and Read, 2020). Recent mechanistic reviews further emphasize keratinocyte-intrinsic metabolic, barrier and transcriptional programs as major determinants of lesion maintenance (Griffiths et al., 2021; Zhou et al., 2022).

Per- and polyfluoroalkyl substances (PFAS) are synthetic fluorinated chemicals characterized by environmental persistence, widespread human exposure and diverse toxicological effects. Legacy and replacement PFAS have been associated with immune, metabolic, endocrine, developmental and hepatic perturbations, although effects vary by congener, exposure context and analytical design (Buck et al., 2011; Sunderland et al., 2019). Human exposure typically occurs as correlated mixtures rather than isolated compounds, and long biological persistence makes sulfonated PFAS especially relevant for chronic internal exposure (Fenton et al., 2021). Perfluorooctanesulfonic acid (PFOS) and perfluorohexanesulfonic acid (PFHxS) are perfluoroalkyl sulfonate (PFSA) chemicals with structural read-across and toxicological relevance. PFHxS is particularly persistent and has been implicated in lipid and immune-related perturbations, including peroxisome proliferator-activated receptor (PPAR)-related mechanisms (Olsen et al., 2007; He et al., 2024).

The skin is a direct environmental interface and an immunologically active epithelial organ. Recent experimental work suggests that PFAS can interact with skin-barrier and immune processes, but the molecular relationship between PFOS- and PFHxS-responsive programs and psoriasis-associated keratinocyte states remains insufficiently characterized (Ragnarsdóttir et al., 2024; Weatherly et al., 2024). Conventional network toxicology can nominate chemical-target or disease-gene intersections, but such analyses may be vulnerable to weak compound specificity when exposure-induced transcriptomic evidence is absent. Conversely, single-cell and spatial psoriasis atlases provide cellular and tissue localization but do not directly identify environmental-response programs. An integrative framework combining chemical prioritization, exposure transcriptomics, disease transcriptomics, single-cell mapping, spatial validation and virtual perturbation can therefore provide a stronger hypothesis-generating bridge between exposure and disease biology.

In this study, we constructed a revised computational toxicology workflow focused on PFOS- and PFHxS-responsive psoriasis programs. We first prioritized six PFAS using chemical structure, evidence layers, exposure differentially expressed gene (DEG) burden, adverse outcome pathway (AOP)-like key-event coverage and read-across analysis. We then extracted PFAS exposure signatures from GSE236956 and compared them with psoriasis bulk transcriptomes from GSE13355. We further mapped the PFOS and PFHxS-associated program to single-cell psoriasis compartments, refined the keratinocyte state structure, assessed candidate regulators and intercellular signaling, evaluated spatial localization in psoriasis tissue sections and performed token-aware Geneformer-compatible virtual perturbation. The goal was not to infer causality from public and computational data alone, but to define a coherent and experimentally testable toxicogenomic hypothesis linking persistent PFSA exposure programs with keratinocyte-centered inflammatory niches in psoriasis.

## 2. Materials and Methods

### 2.1 Study design and public data sources

The study integrated four analytical layers: chemical toxicology prioritization, exposure transcriptomics, psoriasis transcriptomic convergence and tissue-context modeling. Six PFAS were evaluated: perfluorooctanoic acid (PFOA), perfluorohexanoic acid, perfluorobutanoic acid, PFOS, PFHxS and perfluorobutanesulfonic acid. Chemical identifiers and descriptors were checked against PubChem where required (Kim et al., 2023). The exposure transcriptomic layer used the cleaned GSE236956 PFAS differential-expression table, derived from human embryonic stem cell-derived epithelial-lineage models exposed to 10 μM PFAS for 8-16 days depending on lineage and contrast. Because this study reanalyzed public in vitro exposure data rather than applying new chemicals, the dose was interpreted as a controlled hazard-identification condition and not as a direct estimate of human skin target-site concentration. The psoriasis disease transcriptomic layer used GSE13355 as the primary bulk psoriasis expression signature. Single-cell analysis used GSE151177 (Kim et al., 2021), and spatial transcriptomic analysis used GSE202011. All datasets were publicly available and de-identified; no new human samples were collected for this study.

The analysis proceeded from compound-level evidence to cell- and tissue-level localization. First, a read-across and evidence-layer prioritization framework was used to rank the six PFAS for skin-relevant toxicogenomic investigation. Second, PFAS differential-expression signatures were summarized at the genome-wide, gene-set and curated toxicological-module levels. Third, PFOS, PFHxS and PFSA-mean signatures were compared with psoriasis bulk transcriptomes. Fourth, the resulting PFOS and PFHxS-associated keratinocyte program was localized in single-cell and spatial tissue contexts. Finally, candidate regulators, ligands and effector genes were prioritized using regulator-activity scoring, NicheNet/CellChat-style communication analysis, principal component analysis (PCA) virtual-cell perturbation and token-aware Geneformer-compatible perturbation.

### 2.2 Computational toxicology prioritization and read-across scoring

For chemical-level prioritization, scaled descriptor matrices were assembled for each PFAS using chain length, molecular weight, number of fluorine atoms, sulfonate status and skin-prior evidence. Molecular fingerprints were compared using Tanimoto similarity, and Tanimoto-distance-based chemical-space embedding was used to visualize sulfonate and carboxylate group structure. Toxicology evidence layers were curated across the Comparative Toxicogenomics Database (CTD), the CompTox Chemicals Dashboard (CompTox), Toxicity Forecaster evidence, skin/inflammation relevance, PFAS regulatory relevance, data availability and AOP-like key-event coverage (Davis et al., 2023; Richard et al., 2016; Williams et al., 2017). Key-event coverage summarized xenobiotic response, oxidative stress, nuclear factor-κB (NF-κB)/tumor necrosis factor-α (TNF-α) signaling, Janus kinase (JAK)/signal transducer and activator of transcription (STAT) signaling, lipid/PPAR biology, keratinocyte stress, barrier biology and immune programs.

An integrated toxicology prioritization score was calculated as a scaled multi-layer composite of structure, database evidence, transcriptomic burden and skin/AOP support. The score was used to rank PFAS for downstream exposure-disease convergence analyses. The chemical-gene network linking PFOS, PFHxS and PFOA to skin/inflammatory keratinocyte genes was treated as an evidence-prior visualization, not as proof of direct experimentally validated chemical-gene causality. Read-across interpretation emphasized whether a compound combined structural similarity to the PFOS and PFHxS axis with transcriptomic and skin/AOP support.

### 2.3 PFAS exposure signature analysis

Genome-wide PFAS exposure responses were derived from the cleaned GSE236956 differential-expression table. For each PFAS, log2 fold-change (log2FC) values were compared across shared genes using Spearman correlation. Significant DEGs were defined using false discovery rate (FDR) < 0.05 and absolute log2FC ≥ 1.0, and the upregulated and downregulated burdens were visualized across compounds. A PFHxS-specific volcano plot was generated to highlight biologically interpretable genes while avoiding over-labeling of poorly annotated transcripts.

Curated toxicogenomic candidate genes were evaluated across the six PFAS by dot plots in which color represented log2FC and dot size represented -log10(FDR). Candidate genes included inflammatory keratinocyte genes, keratinization and stress markers, cytoskeletal genes, extracellular matrix/transforming growth factor-β (TGF-β) genes, developmental genes and lipid/PPAR-related genes. PFOS-versus-PFHxS contrastive scatter analysis was used to evaluate directionality and strength of shared transcriptional responses. Toxicological module footprints were calculated as the mean log2FC of genes within curated modules under each PFAS exposure.

### 2.4 Pathway, regulator and PFSA footprint analysis

Pathway-level analysis summarized Hallmark pathway activity across PFAS exposure contrasts using normalized enrichment scores (NESs) based on curated Hallmark gene sets (Liberzon et al., 2015). Emphasized pathways included TNF-α/NF-κB, interferon-γ, interleukin (IL)-6/JAK/STAT3, inflammation, p53, apoptosis, hypoxia, unfolded-protein response, epithelial-mesenchymal transition, KRAS up, complement and estrogen early response. PFOS and PFHxS pathway concordance was evaluated by scatter analysis of pathway NES values and categorized into immune/inflammation, remodeling, stress/damage and hormone/other groups.

A pathway/regulator footprint matrix was constructed using curated pathway and transcription-factor modules. Pathway modules included NF-κB, TNF-α, JAK/STAT, mitogen-activated protein kinase/activator protein 1 (AP-1), p53, TGF-β, hypoxia, epidermal growth factor receptor, phosphoinositide 3-kinase-protein kinase B, Wnt, vascular endothelial growth factor and estrogen. Candidate regulators included RELA/NF-κB, STAT1, STAT3, JUN/FOS, aryl hydrocarbon receptor (AHR), PPAR, TP53, suppressor of mothers against decapentaplegic homolog (SMAD), hypoxia-inducible factor 1α (HIF1A), interferon regulatory factor and CEBPB. PFOS-only, PFHxS-only, shared and discordant gene categories were then quantified to distinguish broad PFOS-dominant exposure effects from smaller but directionally informative PFHxS and shared PFSA components.

### 2.5 Psoriasis transcriptomic convergence analysis

GSE13355 was used as the primary psoriasis bulk transcriptomic signature. Differential expression between psoriasis and control samples generated a ranked disease signature. Rank-level concordance between psoriasis log2FC and PFOS, PFHxS, PFSA mean and PFSA rank signals was assessed using Spearman correlation. Gene-level convergence plots compared PFOS, PFHxS and PFSA mean log2FC values with psoriasis log2FC values across shared genes, highlighting high-ranking convergent genes.

Module-level convergence was evaluated by calculating mean log2FC values for toxicological modules across PFOS, PFHxS, PFSA mean and psoriasis. PFSA-psoriasis program genes were visualized using dot plots in which dot color represented log2FC and dot size represented a convergence score. Top convergent genes were ranked using an integrated PFSA-psoriasis convergence score and displayed as a lollipop plot. These analyses were designed to define an exposure-disease bridge for downstream single-cell and spatial evaluation.

### 2.6 Single-cell RNA sequencing atlas, MiloR/Augur analysis and keratinocyte-state refinement

Single-cell RNA sequencing data were processed using standard quality-control, normalization, dimensionality-reduction and clustering workflows, and cell-state structure was visualized using uniform manifold approximation and projection (UMAP) (McInnes et al., 2018). Clusters were annotated using canonical marker genes and mapped to short labels representing activated keratinocyte/PFAS-associated states, basal keratinocytes, T cell/natural killer cells, myeloid cells, melanocytes and fibroblast/PFAS-associated states. A PFOS/PFHxS module score was calculated at single-cell resolution using genes selected from the exposure-disease convergence and toxicological-module analyses. High-score cells were defined using score thresholds and summarized by cluster, disease group and sample.

MiloR neighborhood abundance testing and Augur cell-type prioritization were retained to assess whether PFOS/PFHxS-associated scores aligned with disease-shifted neighborhoods and cell types that best discriminated disease status. For keratinocyte-state refinement, keratinocyte-like cells were subset and annotated into basal, differentiated, activated and cycling states based on marker-gene expression. Marker-derived activation trajectories were calculated, and PFOS/PFHxS scores were compared across states, disease groups and trajectory bins. Trajectory-binned module heatmaps summarized PFOS/PFHxS, basal, differentiation, inflammatory, cycling and stress/AP-1 module activity.

### 2.7 Regulatory and intercellular support analyses

Candidate upstream regulators were evaluated using a regulator-activity and evidence-scoring framework. Regulator activity was summarized across keratinocyte states, and evidence scores incorporated state association, program activity, target support and overall evidence. Regulators included CEBPB, AP-1, NF-κB, STAT, PPAR/lipid, HIF1A, SMAD/TGF-β and AHR. Regulator-target matrices were generated to visualize the relationship between candidate regulators and PFOS/PFHxS-associated inflammatory keratinocyte genes.

NicheNet-style ligand-target analysis was used to prioritize candidate ligands predicted to regulate PFOS/PFHxS-associated keratinocyte target genes. Ligand activity rankings, ligand-target regulatory-potential heatmaps and ligand-target link maps were generated. CellChat-style communication analysis summarized global communication patterns, keratinocyte-directed interactions and pathway-level information-flow patterns.

### 2.8 Spatial transcriptomic validation and niche modeling

Spatial transcriptomic data were processed at the spot level using normalized expression matrices and harmonized spatial coordinates. PFOS/PFHxS core, inflammatory keratinocyte, epidermal stress and keratinocyte module scores were calculated for each spot and projected onto tissue sections. Sample-level distributions were summarized using mean spot scores. Tangram-style transfer was used to estimate keratinocyte-state enrichment in spatial spots, including relative enrichment of PFOS/PFHxS-positive keratinocyte-like states.

Spatial associations were evaluated using sample-level correlation heatmaps, spot-level Spearman correlations and paired low-versus-high niche comparisons. Co-high spot burden was calculated to summarize regions with simultaneous elevation of PFOS/PFHxS score, inflammatory keratinocyte score and/or keratinocyte-state enrichment. A multiview intercellular spatial modeling (MISTy)-style predictor model was used to rank the relative importance of PFOS/PFHxS core, epidermal stress, keratinocyte, inflammatory keratinocyte, myeloid, differentiation, barrier, fibroblast, endothelial and T-cell modules in explaining spatial niche structure. A sample-level module heatmap summarized cross-section heterogeneity.

### 2.9 Virtual-cell and token-aware Geneformer-compatible effector prioritization

Candidate effector genes were prioritized using a dual-mode virtual perturbation framework. The original PCA virtual-cell framework evaluated whether virtual deletion or overexpression shifted cells away from or toward the PFOS/PFHxS-associated state. Deletion rescue, PFOS/PFHxS reduction proxy, overexpression activation and inflammatory/stress module shifts were summarized for candidate genes.

Token-aware Geneformer-compatible in silico perturbation was incorporated as an additional representation-level evidence layer. Geneformer overexpression was interpreted as broad activation-mode evidence. Geneformer deletion was interpreted only when the target gene token was present in the input rank-value sequence, because deletion is meaningful only for token-available cells. Weighted Geneformer support was calculated as 0.70 × Geneformer overexpression activation + 0.30 × Geneformer deletion rescue × token confidence. The final effector score combined original integrated evidence, token-aware Geneformer support and bootstrap stability using the formula: final score = 0.72 × original integrated score + 0.20 × token-aware Geneformer support + 0.08 × bootstrap stability.

### 2.10 Statistical analysis and reproducibility

Unless otherwise specified, gene-level correlations were assessed using Spearman rank correlation. Differential-expression thresholds used FDR < 0.05 and absolute log2FC ≥ 1.0 where binary DEG calls were required. FDR correction followed the Benjamini-Hochberg procedure (Benjamini and Hochberg, 1995). Module scores were calculated from normalized expression values by averaging genes within curated signatures after scaling where appropriate. Group comparisons in single-cell and spatial analyses were interpreted using effect direction, consistency across modules and biological plausibility rather than relying solely on nominal P values, because large cell and spot counts can produce statistically significant but biologically small effects.

Analyses were performed using R and Python workflows for differential expression, enrichment analysis, single-cell processing, spatial processing, communication analysis, virtual perturbation and visualization. Major methods and software frameworks included limma, gene set enrichment analysis, Seurat, Scanpy, Augur, MiloR, NicheNet, CellChat, Tangram, MISTy, Geneformer and ggplot2 (Ritchie et al., 2015; Subramanian et al., 2005; Satija et al., 2015; Stuart et al., 2019; Wolf et al., 2018; Skinnider et al., 2021; Dann et al., 2022; Browaeys et al., 2020; Jin et al., 2021; Biancalani et al., 2021; Tanevski et al., 2022; Theodoris et al., 2023; Wickham, 2016).

## 3. Results

### 3.1 Multi-layer prioritization identified the PFOS and PFHxS axis as the strongest skin-relevant toxicogenomic signal

We first evaluated six PFAS using a chemical and toxicology evidence-prioritization framework. Descriptor analysis separated sulfonated PFAS from carboxylates by sulfonate status, chain length, molecular weight, fluorine count and skin-prior features (Fig. 1A). Molecular fingerprint analysis showed high Tanimoto similarity within the PFAS set (Fig. 1B), and Tanimoto-distance embedding grouped sulfonates and carboxylates into distinct chemical-space regions (Fig. 1C). The toxicology evidence matrix indicated stronger combined support for PFOS and PFHxS than for shorter-chain PFAS across CTD/CompTox, skin/inflammation, regulation, data availability and AOP-like evidence layers (Fig. 1D). AOP-like key-event coverage was also highest for PFOS and PFHxS across xenobiotic, oxidative-stress, NF-κB/TNF, JAK/STAT, lipid/PPAR, keratinocyte-stress, barrier and immune modules (Fig. 1E).

**Figure 1.**
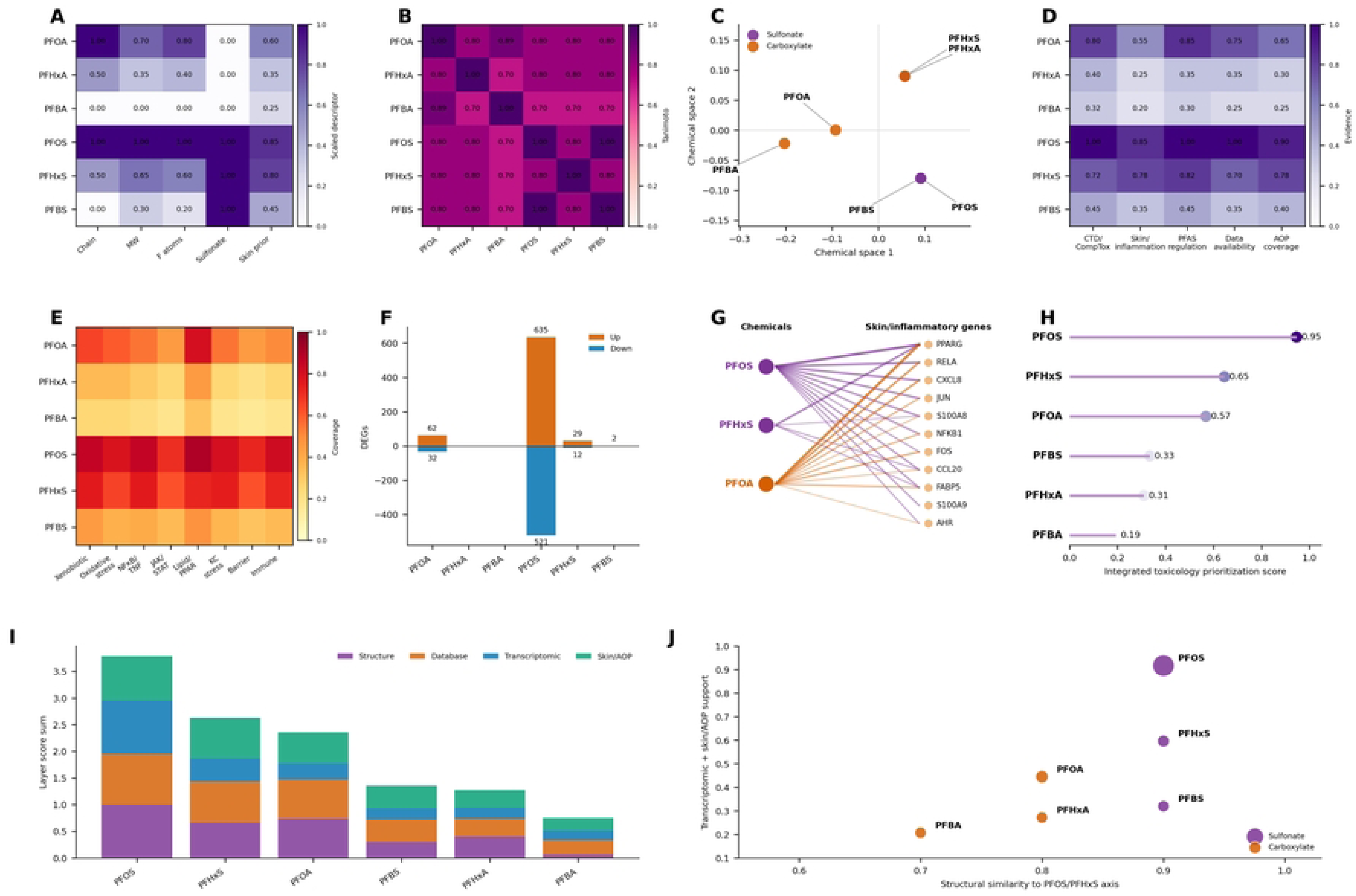
Computational toxicology prioritization of the PFOS/PFHxS axis. (A) Scaled chemical descriptor matrix across six PFAS. (B) Molecular fingerprint/Tanimoto similarity heatmap. (C) Chemical-space embedding based on Tanimoto distance, showing sulfonate and carboxylate separation. (D) Toxicology evidence matrix integrating CTD/CompTox, skin/inflammation, PFAS regulation, data availability and AOP-like support. (E) AOP-like key-event coverage matrix. (F) Exposure DEG burden across PFAS. (G) Chemical-gene evidence-prior network linking PFOS, PFHxS and PFOA to skin/inflammatory keratinocyte genes. (H) Integrated toxicology prioritization score. (I) Evidence-layer composition. (J) Read-across biplot linking structural similarity to the PFOS/PFHxS axis with transcriptomic and skin/AOP support.

Exposure DEG burden further separated PFOS from the remaining PFAS while retaining a smaller PFHxS signal (Fig. 1F). The evidence-prior chemical-gene network linked PFOS, PFHxS and PFOA to inflammatory keratinocyte nodes including PPARG, RELA, CXCL8, JUN, S100A8, NFKB1, FOS, CCL20, FABP5, S100A9 and AHR (Fig. 1G). Integrated scoring ranked PFOS highest with a score of 0.95, followed by PFHxS (0.65) and PFOA (0.57), whereas perfluorobutanesulfonic acid, perfluorohexanoic acid and perfluorobutanoic acid showed lower values (Fig. 1H). Evidence-layer composition showed that PFOS combined strong structural, database, transcriptomic and skin/AOP support, while PFHxS retained multi-layer but weaker support (Fig. 1I). The read-across biplot positioned PFOS and PFHxS as the principal compounds linking structural similarity to transcriptomic and skin/AOP evidence (Fig. 1J).

### 3.2 PFHxS produced a smaller but directionally informative exposure signature within a broader PFOS-dominant PFSA response

Genome-wide correlation analysis across GSE236956 exposure contrasts showed positive concordance among all six PFAS signatures (Fig. 2A). PFHxS correlated most strongly with PFOA and perfluorohexanoic acid (Spearman r = 0.68 for both), followed by perfluorobutanoic acid and perfluorobutanesulfonic acid, and correlated with PFOS at r = 0.60. These correlations suggested a shared PFAS-responsive transcriptional backbone, while incomplete concordance indicated compound-dependent response differences.

**Figure 2.**
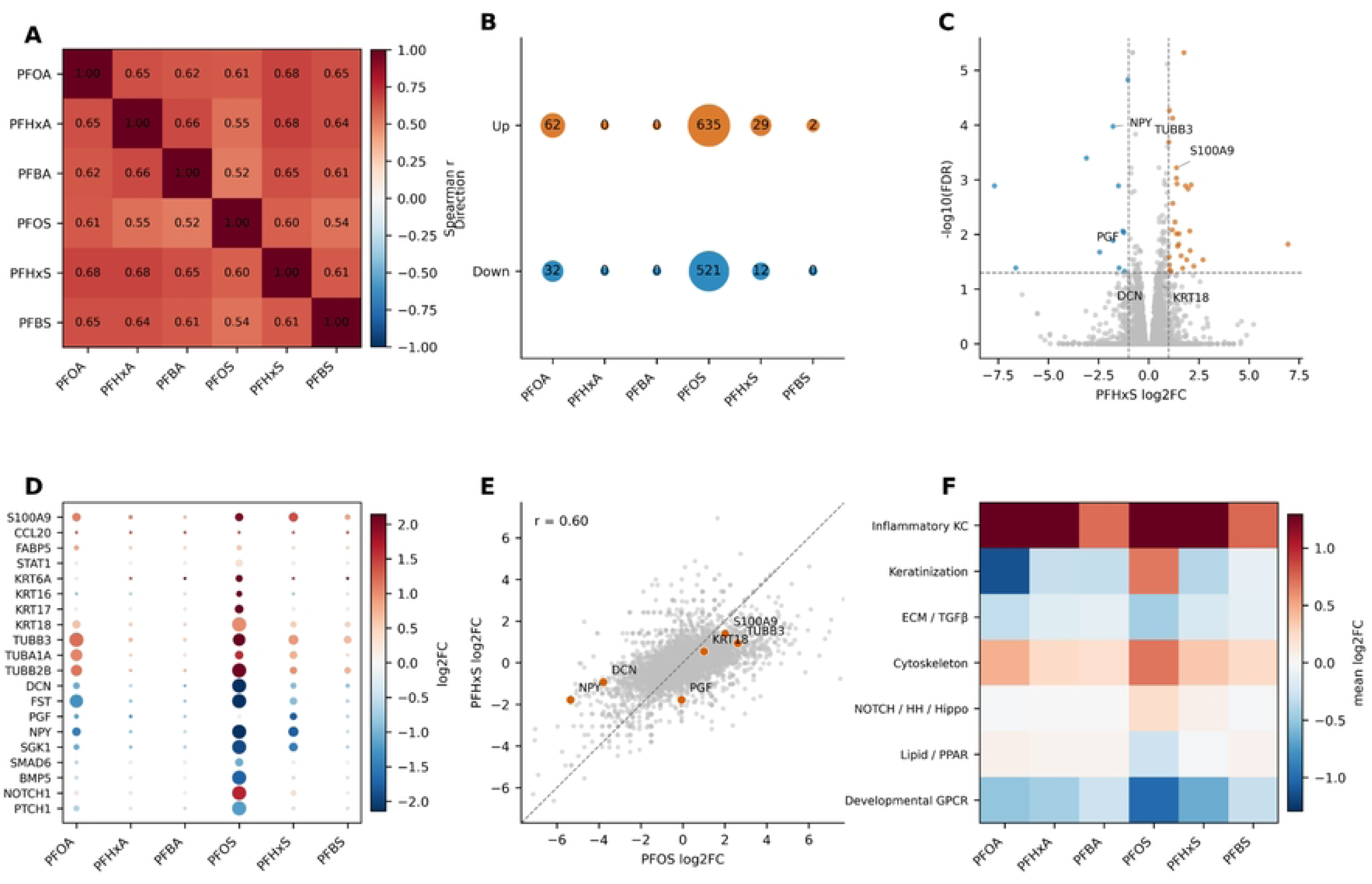
PFAS exposure transcriptomic signature and PFHxS response profile. (A) Genome-wide Spearman correlation heatmap of log2FC values across six PFAS. (B) Bubble matrix of significant upregulated and downregulated DEGs using FDR < 0.05 and absolute log2FC >= 1.0. (C) PFHxS volcano plot with selected biologically interpretable genes labeled. (D) Dot plot of curated toxicogenomic candidate genes across six PFAS; dot color indicates log2FC and dot size indicates -log10(FDR). (E) PFOS versus PFHxS contrastive log2FC scatter. (F) Curated toxicological module footprint represented by mean log2FC for genes in each module under each PFAS exposure.

DEG burden analysis showed that PFOS induced the largest transcriptional response, with 635 upregulated and 521 downregulated genes under the selected threshold; PFOA showed a smaller response with 62 upregulated and 32 downregulated genes, and PFHxS generated 29 upregulated and 12 downregulated genes (Fig. 2B). In the PFHxS volcano plot, S100A9, KRT18 and TUBB3 were among the labeled upregulated genes, whereas DCN, NPY and PGF represented downregulated or context-sensitive genes (Fig. 2C). This pattern supported PFHxS as a lower-amplitude but biologically interpretable exposure signature.

Curated candidate-gene dot plots showed that inflammatory keratinocyte genes, keratinization markers, cytoskeletal genes and developmental or extracellular matrix/TGF-β genes were differentially affected across PFAS (Fig. 2D). The PFOS-versus-PFHxS contrastive scatter showed positive concordance between the two sulfonates (r = 0.60), indicating shared directionality despite stronger PFOS magnitude (Fig. 2E). Module-level footprints suggested that inflammatory keratinocyte and cytoskeletal modules were most responsive to the PFSA axis, while lipid/PPAR, extracellular matrix/TGF-β, NOTCH/Hedgehog/Hippo and developmental G-protein-coupled receptor modules contributed compound-specific signals (Fig. 2F).

### 3.3 Pathway and regulator footprints linked PFSA exposure to inflammatory, stress and remodeling programs

Hallmark pathway footprinting showed enrichment across inflammatory, stress and remodeling pathways, including TNF-α/NF-κB, interferon-γ, IL-6/JAK/STAT3, inflammation, p53, apoptosis, hypoxia, unfolded-protein response, epithelial-mesenchymal transition, KRAS up, complement and estrogen early-response programs (Fig. 3A). Direct comparison of PFOS and PFHxS pathway NES values yielded modest positive concordance (r = 0.30), indicating shared biological directionality with substantial compound-specific response strength (Fig. 3B).

**Figure 3.**
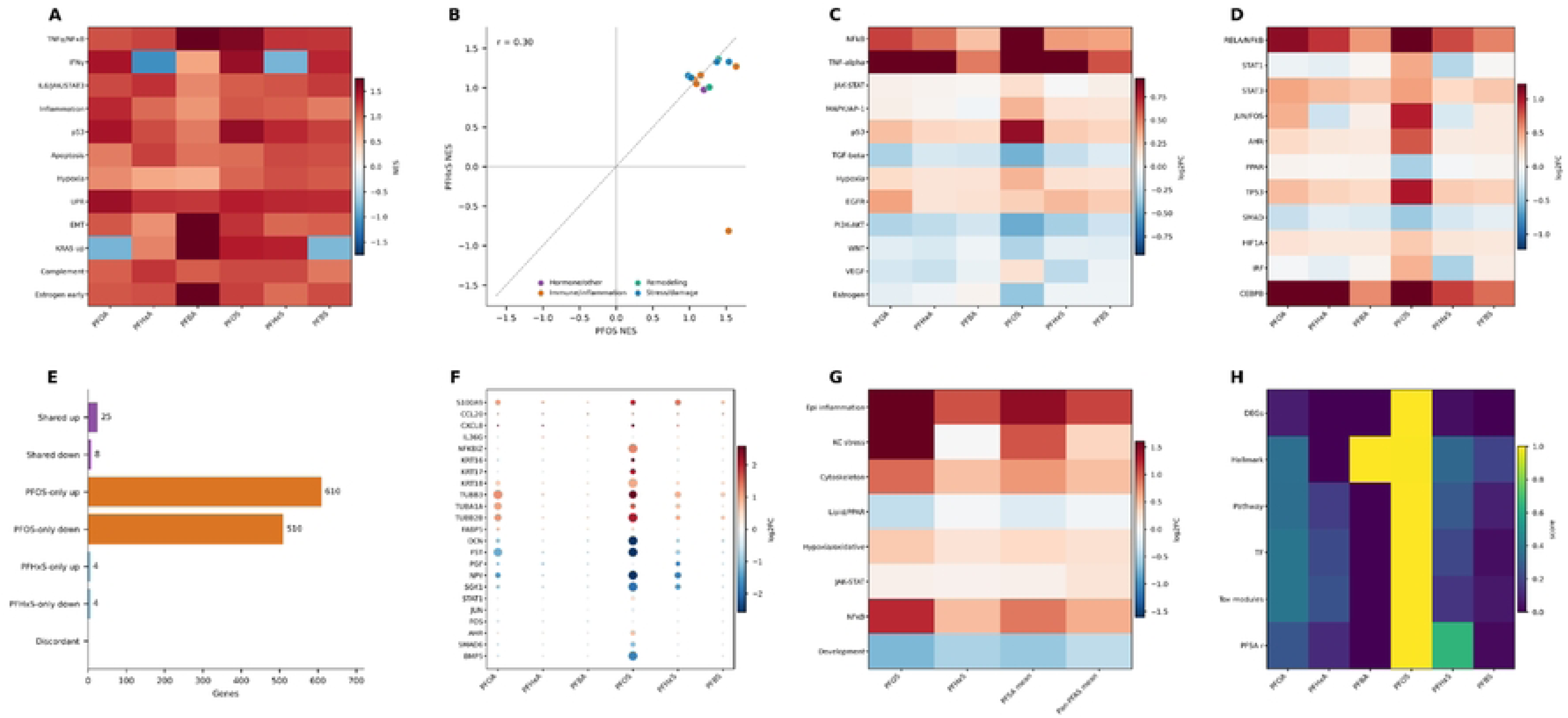
Pathway and regulator footprint of PFAS exposure signatures. (A) Hallmark pathway normalized enrichment score heatmap across six PFAS. (B) PFOS versus PFHxS pathway NES concordance scatter, with pathways grouped into hormone/other, immune/inflammation, remodeling and stress/damage categories. (C) Curated pathway module footprint across PFAS. (D) Candidate transcription-factor/regulator footprint across PFAS. (E) PFOS-only, PFHxS-only, shared and discordant gene categories. (F) Curated candidate-gene dot plot across six PFAS. (G) PFOS, PFHxS, PFSA mean and pan-PFAS mean module footprint. (H) Integrated score heatmap across DEG burden, Hallmark, pathway, transcription-factor, toxicological-module and PFSA-correlation evidence layers.

The curated pathway module footprint further implicated NF-κB, TNF-α, JAK/STAT, mitogen-activated protein kinase/AP-1, p53 and CEBPB-related axes (Fig. 3C). Candidate transcription-factor footprints showed high activity for RELA/NF-κB, STAT1/STAT3, JUN/FOS and CEBPB across PFOS/PFHxS-associated responses, with additional context from AHR, PPAR, TP53, SMAD, HIF1A and interferon regulatory factor (Fig. 3D). Gene-category analysis emphasized the PFOS-dominant nature of the response: PFOS-only upregulated and downregulated categories contained 610 and 510 genes, respectively, whereas shared upregulated and downregulated categories contained 25 and 8 genes and PFHxS-only categories were smaller (Fig. 3E).

Curated toxicological genes including S100A9, CCL20, CXCL8, IL36G, NFKBIZ, KRT16, KRT17, KRT18, TUBB3, FABP5, STAT1, JUN, FOS, AHR, SMAD6 and BMP5 showed differential responses across PFAS and helped define downstream modules (Fig. 3F). Module-level summaries across PFOS, PFHxS, PFSA mean and pan-PFAS mean showed that epithelial inflammation, keratinocyte stress, cytoskeletal remodeling, lipid/PPAR, hypoxia/oxidative, JAK/STAT, NF-κB and developmental modules contributed to the shared footprint (Fig. 3G). Integrated scoring across DEG burden, Hallmark pathways, curated pathways, transcription-factor footprints, toxicological modules and PFSA correlation again prioritized PFOS, with PFHxS contributing a weaker but relevant PFSA-concordant signal (Fig. 3H).

### 3.4 PFSA-responsive programs converged with psoriasis transcriptomic remodeling

We next evaluated whether PFOS/PFHxS-responsive signatures overlapped with psoriasis transcriptomic remodeling in GSE13355. The psoriasis differential-expression table contained 21,653 genes, of which 436 met the selected significance criterion (Fig. 4A). Rank-level concordance between psoriasis and exposure signatures was positive but modest, with Spearman r values of 0.13 for PFOS, 0.11 for PFHxS, 0.13 for PFSA mean and 0.13 for PFSA rank (Fig. 4B). These values indicated that PFSA exposure and psoriasis were not globally identical transcriptomic states, but shared a weak yet consistent disease-relevant direction across the ranked gene space.

**Figure 4.**
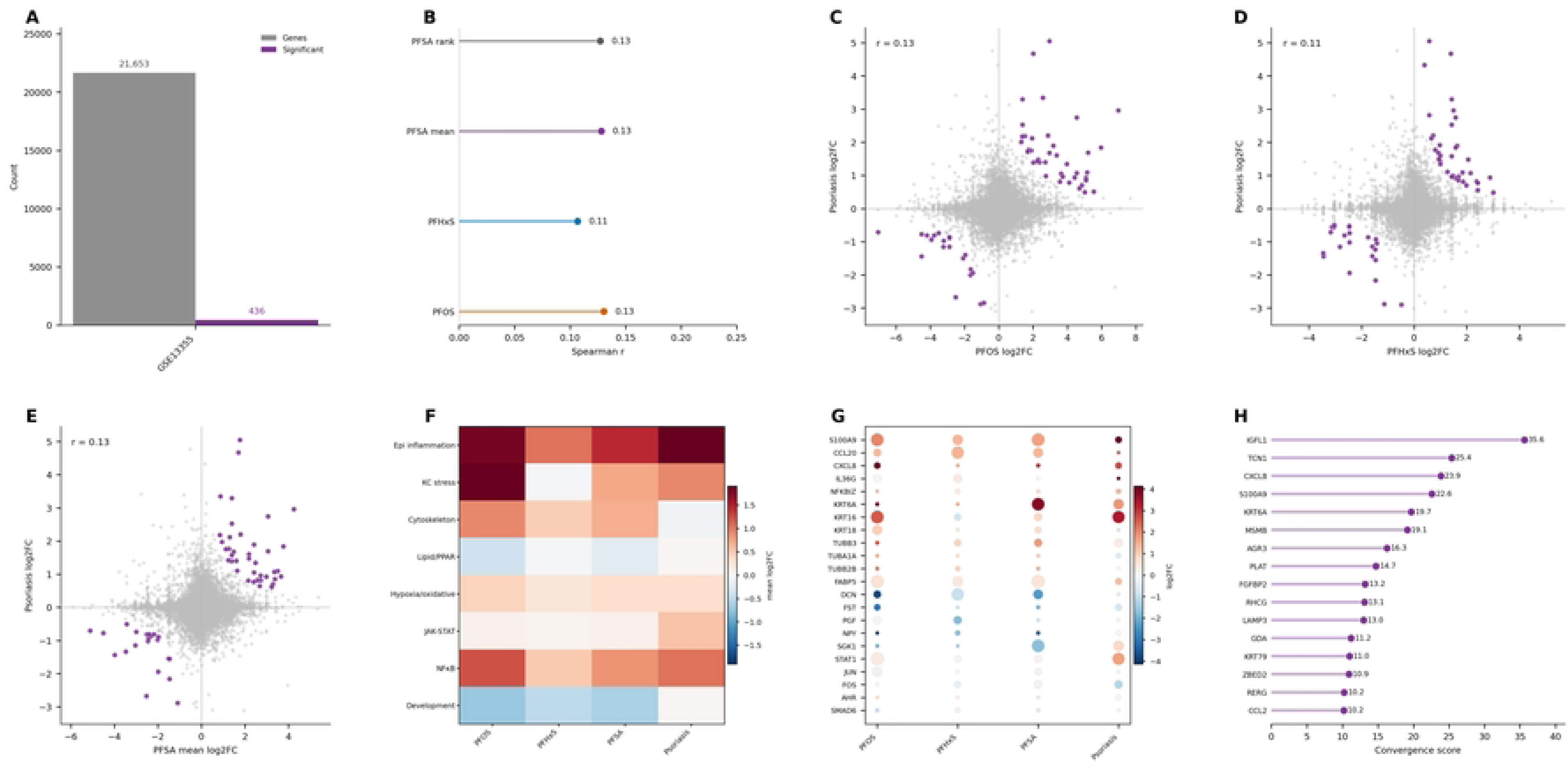
Transcriptomic convergence between PFSA exposure signatures and psoriasis. (A) Psoriasis signature overview from GSE13355 showing total genes and significant genes. (B) Rank- level concordance between psoriasis log2FC and PFOS, PFHxS, PFSA mean or PFSA rank signals using Spearman correlation. (C) PFOS-psoriasis gene-level convergence. (D) PFHxS-psoriasis gene-level convergence. (E) PFSA mean-psoriasis gene-level convergence. Purple points indicate high-ranking convergent genes. (F) Module-level convergence across PFOS, PFHxS, PFSA and psoriasis. (G) PFSA-psoriasis program genes; dot color represents log2FC and dot size represents convergence score. (H) Top convergent genes ranked by PFSA-psoriasis convergence score.

Gene-level convergence plots identified high-ranking overlapping genes in the PFOS-psoriasis comparison (Fig. 4C), the PFHxS-psoriasis comparison (Fig. 4D) and the PFSA mean-psoriasis comparison (Fig. 4E). PFOS showed stronger exposure-associated convergence than PFHxS, consistent with its greater DEG burden. However, PFHxS contributed a directionally concordant component, and the PFSA mean signal captured shared inflammatory and stress-related features.

Module-level convergence showed that epithelial inflammation, keratinocyte stress, cytoskeletal remodeling, lipid/PPAR-related activity, hypoxia/oxidative responses, JAK/STAT, NF-κB and developmental modules were jointly informative across PFOS, PFHxS, PFSA mean and psoriasis (Fig. 4F). PFSA-psoriasis program genes included CXCL8, S100A9, KRT6A and other epidermal activation genes with coordinated log2FC and convergence-score patterns (Fig. 4G). Integrated convergence ranking highlighted IGFL1, TCN1, CXCL8, S100A9, KRT6A, MSMB, AGR3, PLAT, FGFBP2, RHCG, LAMP3, GDA, KRT79, ZBED2, RERG and CCL2 as top convergent genes (Fig. 4H).

### 3.5 Single-cell mapping localized the PFOS/PFHxS program to activated keratinocyte and disease-associated immune compartments

Single-cell analysis resolved 24 clusters that were annotated using marker genes into activated keratinocyte/PFAS-associated clusters, basal keratinocytes, T cell/natural killer cell clusters, myeloid clusters, melanocytes and fibroblast/PFAS-associated cells (Fig. 5A). The PFOS/PFHxS module score was not uniformly distributed across the atlas but was enriched in activated keratinocyte/PFAS-associated clusters and selected myeloid or fibroblast-associated compartments (Fig. 5B). Disease-stratified dot summaries showed a higher high-score cell percentage and mean score in psoriasis-associated compartments than in control-dominant compartments (Fig. 5C). Cluster-level score summaries further confirmed higher mean PFOS/PFHxS scores and higher high-score burdens in activated keratinocyte/PFAS-associated clusters than in many immune or non-epithelial clusters (Fig. 5D).

**Figure 5.**
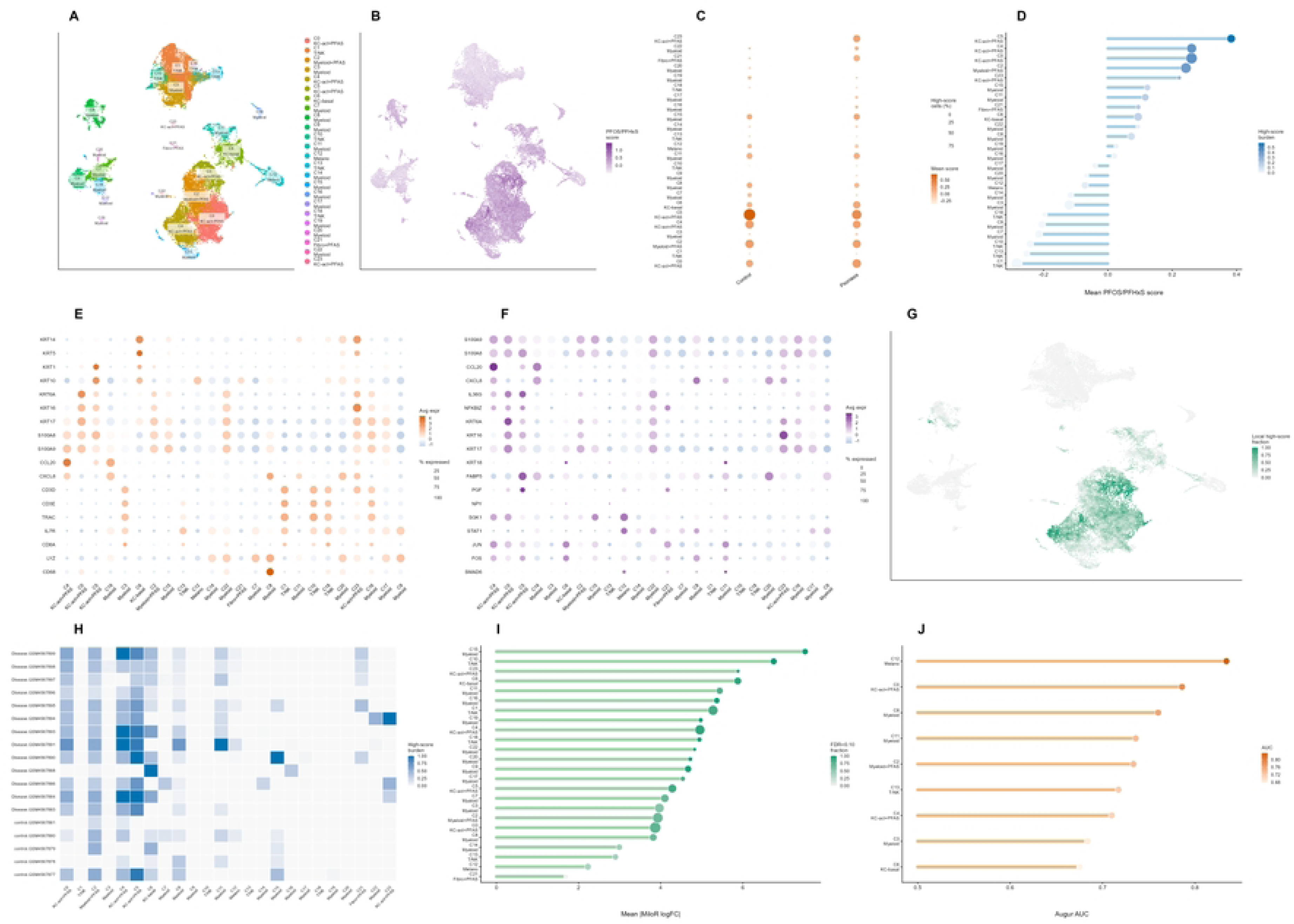
Single-cell atlas mapping of the PFOS/PFHxS-associated program. (A) UMAP visualization of annotated psoriasis skin single-cell clusters. (B) PFOS/PFHxS score projected onto the single-cell UMAP. (C) Dot plot showing PFOS/PFHxS score and high-score cell percentage across control and psoriasis groups. (D) Mean PFOS/PFHxS score and high-score burden across annotated clusters. (E) Dot plot of canonical marker genes for keratinocyte, T cell/natural killer cell and myeloid annotation. (F) Dot plot of PFSA-psoriasis program genes across clusters. (G) Local high-score fraction projected on the single-cell UMAP. (H) Sample-by-cluster high-score burden heatmap. (I) MiloR neighborhood abundance summary. (J) Augur area under the curve prioritization across cell states.

Canonical marker-gene dot plots confirmed the identity of keratinocyte, T cell/natural killer cell and myeloid compartments (Fig. 5E). Dot plots of PFSA-psoriasis program genes showed coordinated expression of S100A9, S100A8, CCL20, CXCL8, IL36G, NFKBIZ, KRT6A, KRT16, KRT17, FABP5, STAT1, JUN, FOS and SMAD6 in disease-relevant cell states (Fig. 5F). Local high-score fraction concentrated in activated keratinocyte-rich regions within the single-cell manifold (Fig. 5G).

Sample-level high-score burden differed across control and disease samples, with disease samples tending to show stronger PFOS/PFHxS-associated cell-state enrichment (Fig. 5H). MiloR neighborhood analysis identified neighborhoods with elevated disease-associated abundance changes (Fig. 5I). Augur prioritized cell states with high disease-discriminating capacity, including basal keratinocyte, myeloid and activated keratinocyte states (Fig. 5J). These results suggested that PFOS/PFHxS-associated signals are embedded in the cellular architecture of psoriatic lesions and are especially connected to keratinocyte activation and myeloid inflammatory contexts.

### 3.6 Keratinocyte-state refinement linked PFOS/PFHxS scores to an activation trajectory

Because the single-cell atlas implicated keratinocytes as a major compartment of the PFOS/PFHxS program, we subset keratinocyte-like cells and annotated four transcriptional states: basal, differentiated, activated and cycling (Fig. 6A). PFOS/PFHxS score projection showed enrichment within activated and partially differentiated regions of the keratinocyte manifold (Fig. 6B). Marker-gene dot plots confirmed state annotation: KRT5, KRT14 and ITGA6 marked basal cells; KRT1, KRT10, IVL and FLG marked differentiation; KRT6A, KRT16, KRT17, S100A8, S100A9, CCL20, CXCL8 and IL36G marked activation; and MKI67/TOP2A marked cycling cells (Fig. 6C).

**Figure 6.**
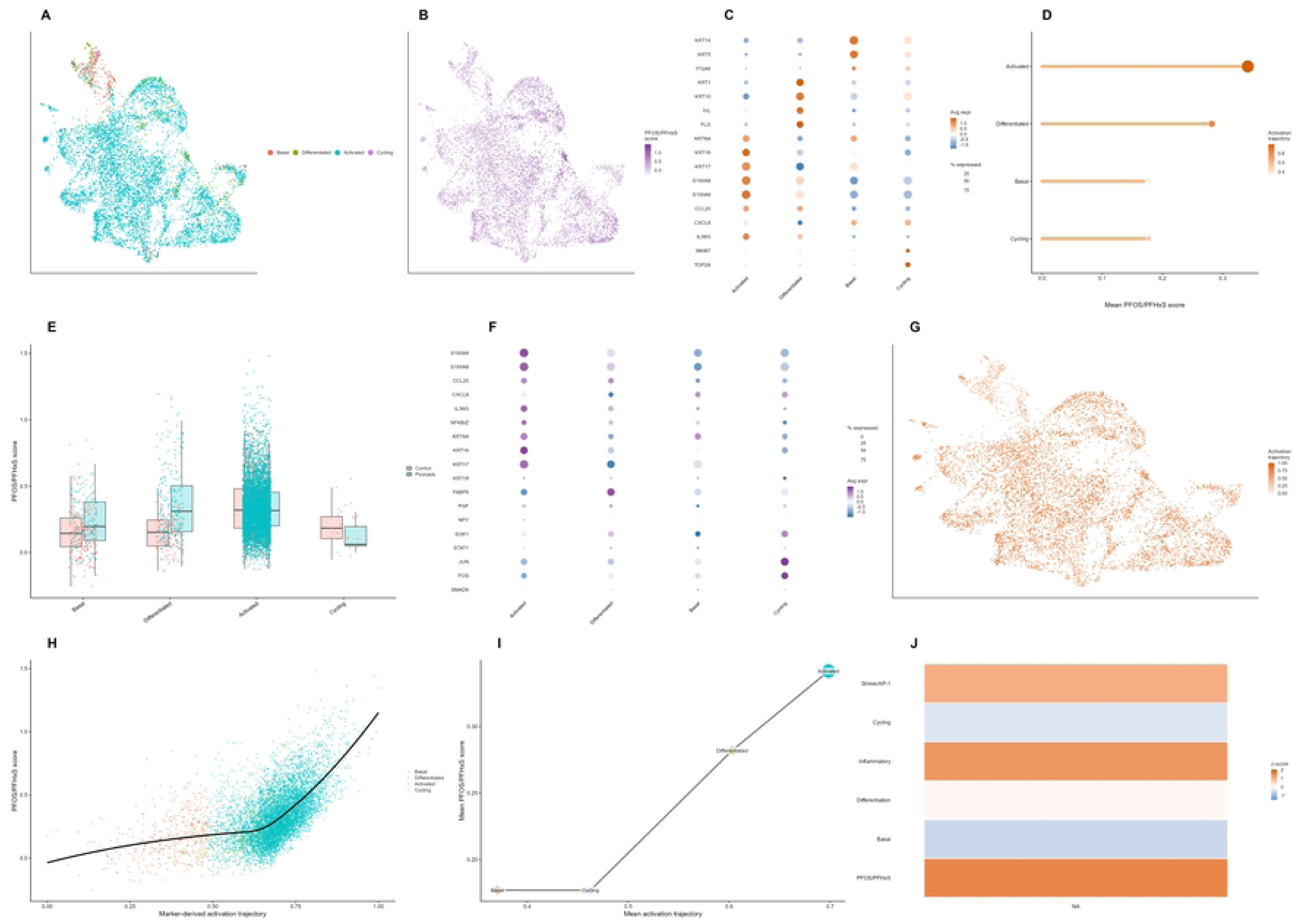
Keratinocyte-state refinement of the PFOS/PFHxS program. (A) UMAP visualization of keratinocyte states, including basal, differentiated, activated and cycling states. (B) PFOS/PFHxS score projected onto keratinocyte UMAP. (C) Dot plot of keratinocyte state marker genes. (D) Mean PFOS/PFHxS score across keratinocyte states. (E) PFOS/PFHxS score distributions by state and disease group. (F) Dot plot of PFSA-psoriasis program genes across keratinocyte states. (G) Marker-derived activation trajectory projected onto UMAP. (H) Relationship between activation trajectory and PFOS/PFHxS score. (I) State-transition graph summarizing mean activation trajectory and mean PFOS/PFHxS score. (J) Trajectory-binned module heatmap.

Mean PFOS/PFHxS scores differed across keratinocyte states and were highest in activated keratinocytes (Fig. 6D). Disease-stratified score distributions suggested that psoriasis keratinocytes were shifted toward higher PFOS/PFHxS scores, especially in activated and differentiated states (Fig. 6E). Program-gene expression patterns supported this interpretation, with S100A9, S100A8, CCL20, CXCL8, IL36G, NFKBIZ, KRT6A, KRT16, KRT17, FABP5, STAT1, JUN and FOS enriched in activated keratinocytes (Fig. 6F).

A marker-derived activation trajectory increased from basal and cycling states toward differentiated and activated states (Fig. 6G). At the single-cell level, PFOS/PFHxS scores rose along this activation trajectory (Fig. 6H). State-level summaries showed a parallel increase in mean activation trajectory and mean PFOS/PFHxS score, supporting an activation-linked exposure-informed program (Fig. 6I). Trajectory-binned module analysis showed high PFOS/PFHxS, inflammatory and stress/AP-1 module activity in activated regions, whereas basal and differentiation modules showed distinct or opposing patterns (Fig. 6J).

### 3.7 Regulatory and intercellular analyses nominated CEBPB, AP-1, NF-κB and inflammatory ligand contexts

We next evaluated candidate regulators and intercellular inputs that could support the PFOS/PFHxS-associated activated keratinocyte state. Regulator-activity heatmaps showed that NF-κB, STAT, AP-1, AHR, PPAR/lipid, CEBPB, HIF1A and SMAD/TGF-β differed across basal, differentiated, activated and cycling keratinocyte states (Fig. 7A). Evidence ranking placed CEBPB, AP-1 and NF-κB at the top, with STAT and PPAR/lipid as secondary modules (Fig. 7B). The regulator-target matrix connected these regulators to downstream inflammatory keratinocyte genes, including S100A9, CCL20, CXCL8 and KRT-family activation markers (Fig. 7C). The evidence matrix further supported CEBPB, AP-1 and NF-κB as high-evidence candidates linked to state specificity, program activity, target-gene support and overall evidence (Fig. 7D).

**Figure 7.**
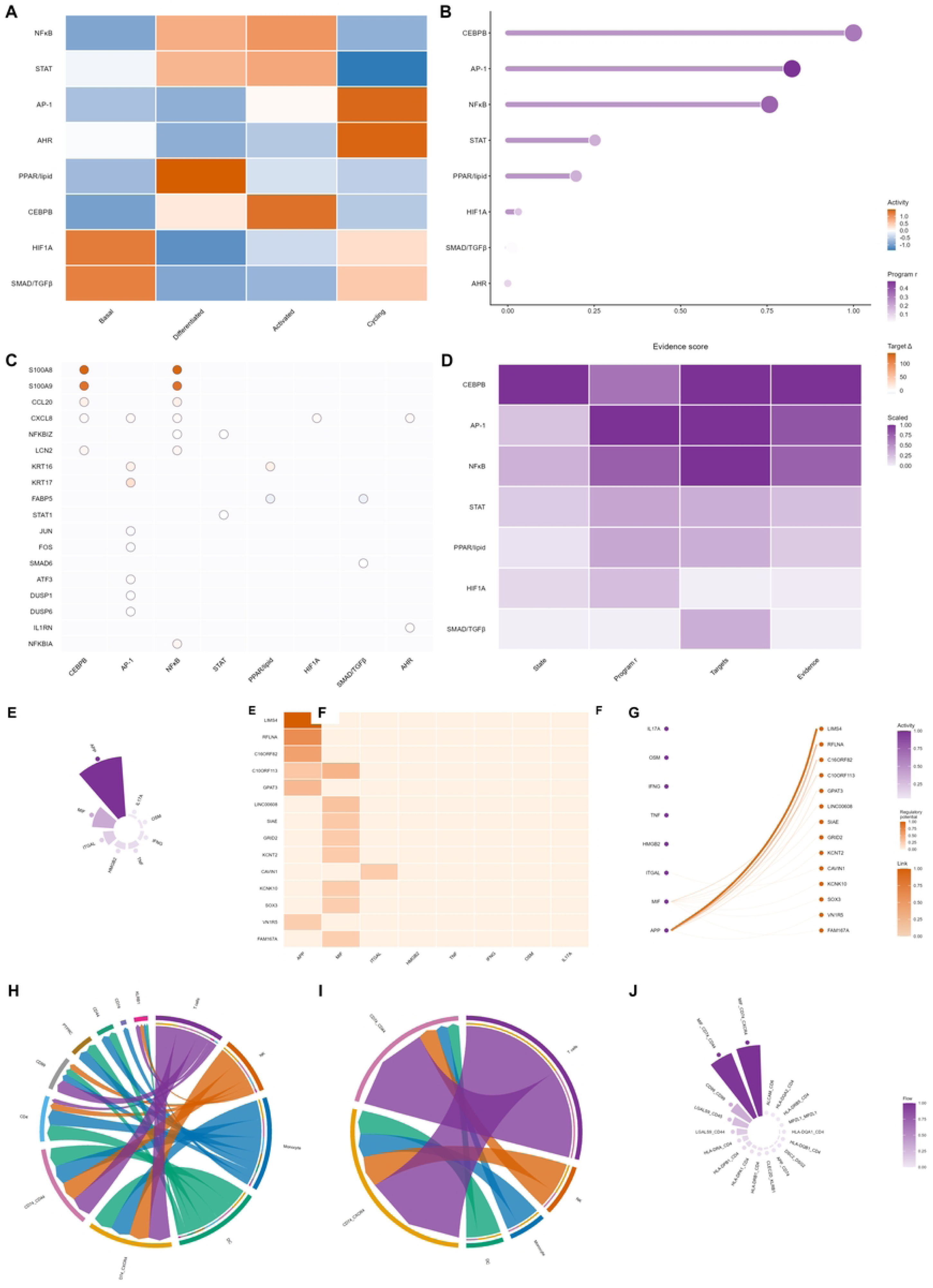
Regulatory and intercellular support for the PFOS/PFHxS-associated keratinocyte program. (A-D) Regulator block showing regulator activity across keratinocyte states, evidence ranking, regulator-target matrix and evidence matrix. (E-G) NicheNet-style ligand-target block showing ligand activity ranking, ligand-target regulatory-potential heatmap and ligand-target link map. (H-J) CellChat-style block showing global communication circle, keratinocyte-directed communication circle and radial information-flow summary. These analyses provide candidate regulatory and microenvironmental support and do not prove causal ligand or regulator perturbation.

NicheNet-style ligand activity analysis nominated candidate inflammatory and alarmin-like ligands predicted to regulate PFOS/PFHxS-associated keratinocyte target genes (Fig. 7E). Ligand-target regulatory-potential heatmaps connected candidate ligands to downstream targets such as S100A9, CCL20, CXCL8, FABP5, KRT16, KRT17 and other inflammatory or stress-associated keratinocyte genes (Fig. 7F). Ligand-target link maps highlighted candidate inputs including IL17A, OSM, IFNG, TNF, HMGB2, ITGAL, MIF and APP-related axes, although these predictions require direct experimental validation (Fig. 7G).

CellChat-style communication analysis showed dense global communication among keratinocyte, T-cell, myeloid, dendritic and stromal compartments (Fig. 7H). Keratinocyte-directed communication emphasized inflammatory and immune-stromal pathways converging on epidermal receiver states (Fig. 7I). Radial information-flow summaries suggested that multiple signaling programs may contribute to the microenvironment of PFOS/PFHxS-associated keratinocyte activation (Fig. 7J). Collectively, the regulator and communication analyses supported a model in which intrinsic keratinocyte inflammatory transcription factors interact with immune and stromal ligand contexts to reinforce the PFOS/PFHxS-associated program.

### 3.8 Spatial transcriptomics validated PFOS/PFHxS-inflammatory keratinocyte niche organization

Spatial transcriptomic mapping showed heterogeneous but organized distribution of PFOS/PFHxS core scores across psoriasis tissue sections (Fig. 8A). Inflammatory keratinocyte scores showed partially overlapping epidermal domains (Fig. 8B), and epidermal stress scores showed a similar but not identical spatial pattern (Fig. 8C). Sample-level module score comparisons indicated elevated mean spot scores in lesional sections for PFOS/PFHxS core, inflammatory keratinocyte, epidermal stress and keratinocyte modules (Fig. 8D). These maps demonstrated that the exposure-informed program was not randomly distributed across tissue but concentrated in spatial domains consistent with keratinocyte-rich inflammatory niches.

**Figure 8.**
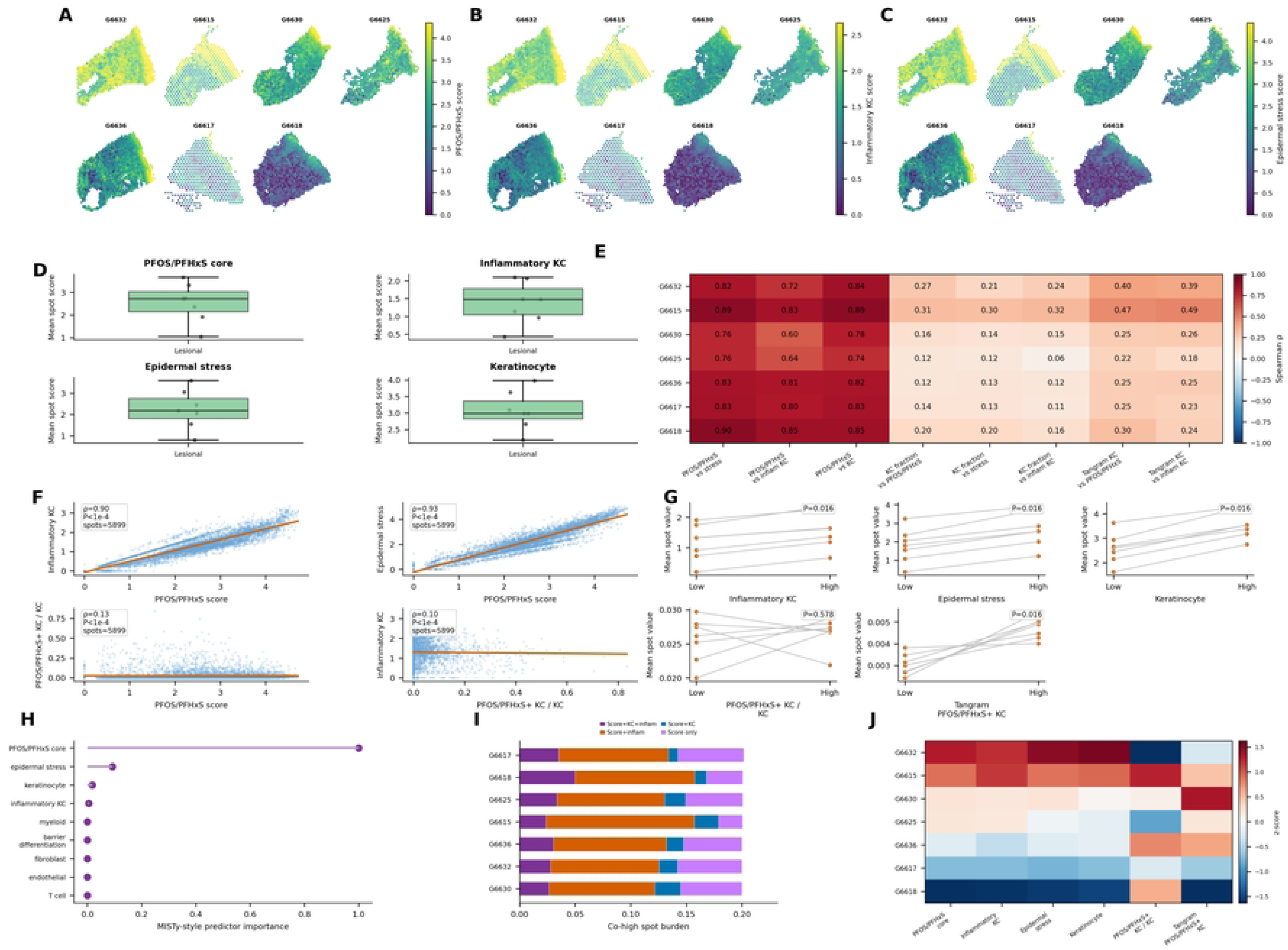
Spatial validation and niche modeling of PFOS/PFHxS-associated programs. (A-C) Spatial score atlases for PFOS/PFHxS core, inflammatory keratinocyte and epidermal stress scores across psoriasis tissue sections. (D) Sample-level module score comparisons. (E) Sample-level correlation heatmap linking PFOS/PFHxS, inflammatory keratinocyte, epidermal stress, keratinocyte and Tangram-derived keratinocyte-state metrics. (F) Spot-level association plots. (G) Paired low-versus-high niche comparisons. (H) MISTy-style predictor importance ranking. (I) Sample-level co-high spot burden. (J) Sample-level module heatmap.

Sample-level correlation heatmaps showed strong positive relationships between PFOS/PFHxS core and inflammatory or stress modules across sections (Fig. 8E). At the spot level, PFOS/PFHxS score was highly correlated with inflammatory keratinocyte score (ρ = 0.90, P < 1 × 10−4) and epidermal stress score (ρ = 0.93, P < 1 × 10−4), whereas correlations involving PFOS/PFHxS-positive keratinocyte fraction were weaker (Fig. 8F). These findings suggested that module-defined PFOS/PFHxS spatial domains capture a broad inflammatory epidermal structure beyond a single mapped cell-state fraction.

Paired low-versus-high niche comparisons showed higher inflammatory keratinocyte, epidermal stress, keratinocyte and Tangram PFOS/PFHxS-positive keratinocyte values in high-score regions, with P = 0.016 for several paired comparisons; the PFOS/PFHxS-positive keratinocyte fraction normalized to keratinocyte content did not show a significant paired change (P = 0.578) (Fig. 8G). MISTy-style predictor analysis ranked the PFOS/PFHxS core score as the dominant predictor, followed by epidermal stress, keratinocyte and inflammatory keratinocyte modules (Fig. 8H). Co-high burden analysis identified tissue sections with simultaneous elevation of PFOS/PFHxS, inflammatory keratinocyte and keratinocyte features (Fig. 8I). A sample-level module heatmap further revealed cross-section heterogeneity in PFOS/PFHxS-inflammatory niche burden (Fig. 8J).

### 3.9 Virtual-cell and token-aware Geneformer analyses prioritized candidate effector genes

Finally, we used virtual-cell and token-aware Geneformer-compatible perturbation to prioritize candidate effector genes within the PFOS/PFHxS-associated keratinocyte program. Integrated evidence heatmaps combined high-low expression, activated-state expression, PCA virtual deletion, PFOS/PFHxS reduction proxy, PCA overexpression, regulator support, Milo/Augur support, Geneformer deletion rescue, Geneformer overexpression activation and bootstrap stability (Fig. 9A). Keratinocyte UMAP visualization confirmed that the candidate perturbation analysis was anchored in the basal, differentiated, activated and cycling state structure (Fig. 9B). PFOS/PFHxS score projection again highlighted activated keratinocyte regions as the main state context for effector prioritization (Fig. 9C).

**Figure 9.**
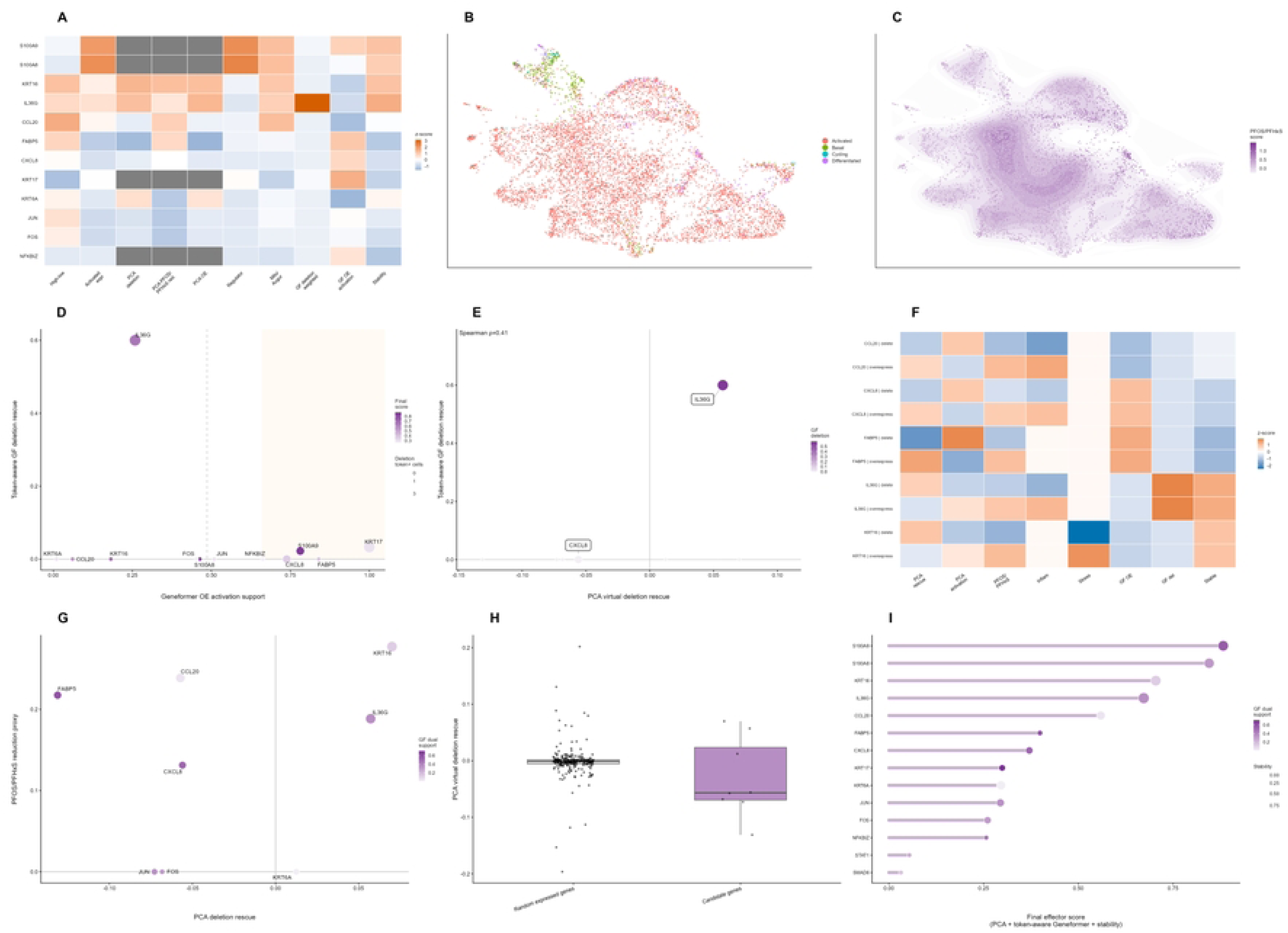
Virtual-cell and token-aware Geneformer-compatible prioritization of candidate effector genes. (A) Integrated candidate-gene evidence heatmap. (B) Keratinocyte UMAP by state. (C) PFOS/PFHxS score projected onto UMAP. (D) Geneformer overexpression activation support versus token-aware Geneformer deletion rescue. (E) Relationship between PCA virtual deletion rescue and token-aware Geneformer deletion rescue. (F) Virtual perturbation heatmap for selected overexpression and deletion operations. (G) PCA deletion rescue versus PFOS/PFHxS reduction proxy, colored by Geneformer dual support. (H) Comparison of PCA virtual deletion rescue between random expressed genes and candidate genes. (I) Final effector score ranking combining PCA virtual-cell evidence, token-aware Geneformer support and bootstrap stability. Geneformer analyses are computational prioritization and not experimental causal proof.

The Geneformer overexpression-versus-deletion scatter showed that broad activation support and token-aware deletion rescue captured complementary perturbation dimensions (Fig. 9D). The relationship between PCA virtual deletion rescue and token-aware Geneformer deletion rescue was positive but moderate (Spearman ρ = 0.41), indicating that the two perturbation frameworks were related but not redundant (Fig. 9E). Perturbation heatmaps showed that deletion and overexpression of CCL20, CXCL8, IL36G, FABP5, KRT16 and KRT17 affected PCA rescue, PCA activation, PFOS/PFHxS, inflammatory, stress, Geneformer overexpression, Geneformer deletion and stability dimensions differently (Fig. 9F).

The PCA deletion-rescue versus PFOS/PFHxS-reduction proxy analysis highlighted genes such as KRT16, IL36G, CCL20, FABP5 and CXCL8 as candidates with distinct rescue and activation profiles (Fig. 9G). Random-gene comparison suggested that candidate genes showed stronger or more structured PCA virtual deletion rescue than randomly selected expressed genes (Fig. 9H). The final effector ranking prioritized S100A9 and S100A8 at the top, followed by KRT16, IL36G, CCL20, CXCL8, FABP5, KRT17, KRT6A, FOS, JUN and NFKBIZ (Fig. 9I). These genes span alarmin biology, keratinocyte activation, chemokine signaling, lipid handling, stress/AP-1 activity and inflammatory transcriptional control.

## 4. Discussion

### 4.1 Principal findings and toxicological framing

This study provides a revised computational toxicology framework linking PFOS-and PFHxS-responsive programs to psoriasis-associated keratinocyte niches. Multi-layer prioritization first identified PFOS as the strongest signal and PFHxS as a weaker but consistent sulfonate read-across component. This interpretation is consistent with the broader PFAS toxicology literature, in which immune, metabolic and epithelial effects vary by compound and exposure context (Fenton et al., 2021). Because human PFAS exposure usually occurs through correlated mixtures, a program-level interpretation is more appropriate than attributing all downstream biology to a single congener (Sunderland et al., 2019).

The main contribution of this work is the convergence of independent evidence layers. Chemical descriptors and evidence scoring nominated the PFOS and PFHxS axis; exposure transcriptomics identified a PFSA-responsive program; psoriasis transcriptomics demonstrated disease convergence; single-cell analysis localized the program to activated keratinocyte and inflammatory compartments; spatial transcriptomics showed tissue organization; and virtual perturbation prioritized testable effectors. These findings do not establish causality, but they provide a coherent mechanistic hypothesis that can be evaluated in skin-relevant toxicology models.

### 4.2 PFOS and PFHxS read-across and exposure-disease convergence

PFOS and PFHxS share sulfonate chemistry and structural features that support read-across, but they differed in response magnitude in our transcriptomic analysis. PFOS produced the strongest DEG burden and integrated prioritization score, whereas PFHxS produced a smaller but directionally informative signature. This distinction is important because PFHxS has a long biological persistence in humans, making even lower-amplitude molecular effects potentially relevant for chronic internal exposure (Olsen et al., 2007). PFAS terminology, classification and environmental persistence further support treating sulfonated PFAS as related but not interchangeable compounds (Buck et al., 2011).

The PFOS and PFHxS axis is biologically plausible for psoriasis-relevant toxicology because psoriasis lesions are characterized by inflammation, metabolic rewiring, endoplasmic reticulum stress, epithelial differentiation defects and barrier remodeling. PFHxS has been linked to lipid homeostasis and PPARα-related mechanisms in experimental systems (He et al., 2024). In addition, PFAS can cross reconstructed human skin barriers, and topical PFHxS exposure can induce local and systemic immune alterations in mice (Ragnarsdóttir et al., 2024; Weatherly et al., 2024). Therefore, the observed overlap with NF-κB/TNF-α, JAK/STAT, AP-1, unfolded-protein response, cytoskeletal and lipid/PPAR modules is mechanistically plausible rather than a purely statistical coincidence.

### 4.3 Keratinocyte inflammatory niches and regulatory context

The keratinocyte-centered findings align with current psoriasis biology. Keratinocytes are not passive structural cells; they produce cytokines, chemokines, antimicrobial peptides and alarmins that sustain lesion inflammation (Nestle et al., 2009). Clinical and translational reviews similarly describe psoriasis as a coordinated epithelial-immune disease involving epidermal activation, immune recruitment and vascular remodeling (Griffiths et al., 2021). In this context, the enrichment of PFOS/PFHxS scores in activated keratinocytes suggests that the exposure-informed program maps to an established disease-relevant cell state.

Several prioritized genes have direct links to psoriatic keratinocyte biology. IL-17-related stimulation can induce CCL20 in keratinocytes, supporting recruitment of CCR6-positive IL-17-producing cells (Harper et al., 2009). The CCL20-CCR6 axis is also recognized as a psoriasis-relevant chemokine pathway (Furue et al., 2020). S100A9 and S100A8 connect the model to alarmin biology, neutrophil-associated inflammation and psoriasiform disease amplification (Mellor et al., 2022; Silva de Melo et al., 2023). FABP5 provides a lipid-metabolic bridge because keratinocyte FABP5 can promote inflammatory recruitment in psoriasis (Hao et al., 2023).

The regulatory and spatial analyses further strengthened the niche model. CEBPB, AP-1 and NF-κB were nominated as upstream contexts, while STAT and PPAR/lipid programs provided secondary support. Regulon-style frameworks such as SCENIC have demonstrated the value of inferring transcription-factor context from single-cell data, but direct phosphorylation or perturbation assays are still required for validation (Aibar et al., 2017). Cell-cell communication and ligand-target inference are appropriate for prioritizing candidate intercellular axes, but they should be interpreted as computational hypotheses rather than direct measurements of ligand activity (Browaeys et al., 2020; Jin et al., 2021). Spatial mapping is also inferential, yet Tangram-style transfer and spot-level scoring are well suited to localize single-cell-derived programs within tissue architecture (Biancalani et al., 2021). The strong co-localization of PFOS/PFHxS core, inflammatory keratinocyte and epidermal stress scores therefore supports a tissue-level inflammatory niche rather than a single isolated gene effect.

### 4.4 Virtual perturbation, limitations and experimental validation

Virtual perturbation prioritized a compact set of effector genes for experimental follow-up. S100A9 and S100A8 ranked highest, consistent with their expression in inflammatory keratinocyte and myeloid-associated compartments. KRT16 and KRT17 reflect keratinocyte activation and hyperproliferation, IL36G and CCL20 connect to cytokine and chemokine loops, CXCL8 links the program to neutrophil recruitment, and FABP5 provides a lipid-inflammatory bridge. Token-aware Geneformer analysis adds a foundation-model perspective, but such representation-level perturbation should be considered prioritization rather than mechanistic proof (Theodoris et al., 2023).

Several computational components require cautious interpretation. MiloR and Augur can identify disease-shifted neighborhoods and cell types, but they depend on the composition and preprocessing of the single-cell atlas (Dann et al., 2022; Skinnider et al., 2021). MISTy-style analysis can rank spatial predictors, but it does not prove causal communication among tissue compartments (Tanevski et al., 2022). Similarly, the evidence-prior chemical-gene network should be strengthened with exported CTD, CompTox and Toxicity Forecaster evidence tables before final submission, because curated database links and predicted targets are not equivalent to direct PFOS or PFHxS perturbation experiments (Davis et al., 2023; Williams et al., 2017).

The study has several limitations. The exposure transcriptomic dataset was not generated from adult psoriatic keratinocytes or skin tissue, and differences in cell type, developmental state, exposure duration and culture context may influence PFAS response signatures. The psoriasis bulk, single-cell and spatial datasets did not include measured PFOS or PFHxS concentrations from the same samples. Therefore, cross-dataset convergence supports biological plausibility but cannot establish that PFOS or PFHxS caused the observed psoriasis programs. In addition, module scores, Tangram-style transfer, CellChat, NicheNet, MISTy-style modeling and Geneformer-compatible perturbation all depend on modeling assumptions and require experimental validation. Future studies should test the proposed PFOS/PFHxS-keratinocyte niche model using exposure-relevant concentrations in primary human keratinocytes, reconstructed human epidermis and organotypic psoriasis-like skin cultures. Experiments should include PFOS and PFHxS separately and in mixtures, with and without cytokine priming by IL-17A, TNF-α, IL-22 or IL-36. Key endpoints should include target-site PFAS accumulation by liquid chromatography-tandem mass spectrometry, barrier markers, lipidomic remodeling, PPAR activity, oxidative and endoplasmic reticulum stress, KRT16/KRT17 activation, S100A8/S100A9 release, CCL20 and CXCL8 secretion, STAT1/STAT3 and NF-κB/AP- 1 activation and spatial organization in tissue-like models. Perturbation of S100A9, FABP5, KRT17, IL36G, CCL20, CXCL8, AP-1 and NF-κB/STAT pathways would directly test whether the predicted effectors mediate PFAS-amplified keratinocyte inflammation.

## 5. Conclusion

This study integrated chemical read-across, toxicology evidence scoring, PFAS exposure transcriptomics, psoriasis transcriptomic convergence, single-cell and spatial mapping, regulatory inference and token-aware virtual perturbation. The results support a computational toxicogenomic hypothesis that PFOS- and PFHxS-responsive programs intersect with psoriasis-associated keratinocyte inflammation, epidermal stress and spatial niche organization. PFOS showed the strongest signal, whereas PFHxS contributed a smaller but relevant PFSA-concordant component. Candidate effectors, including S100A9, S100A8, KRT16, IL36G, CCL20, CXCL8, FABP5, KRT17, FOS, JUN and NFKBIZ, provide focused targets for future experimental validation in keratinocyte-centered inflammatory skin models.

## CRediT authorship contribution statement

**Junhao Ma**: Conceptualization, Methodology, Formal analysis, Visualization, Writing – original draft. **Junjie Tan**: Methodology, Data curation, Formal analysis. **Yucan Wang**: Data curation, Validation. **Shihao Zhou**: Investigation, Validation. **Shanshan Cai**: Conceptualization, Supervision, Project administration, Writing – review & editing. **Qianying Yu**: Conceptualization, Supervision, Project administration, Writing – review & editing.

## Declarations

This study used only publicly available and de-identified datasets. No new human participants, clinical samples or animal experiments were involved; therefore, additional ethics approval was not required.

## Acknowledgements

The authors gratefully acknowledge the financial support from the School-Hospital Joint Innovation Fund of Chengdu University of Traditional Chinese Medicine [WXLH202403142], the 2025 Undergraduate Innovation Training Program of Chengdu University of Traditional Chinese Medicine [202510633034], and the Student Scientific Research Practice Innovation Project of Chengdu University of Traditional Chinese Medicine [ky-2026041].

## Data Availability

The datasets analyzed in this study are publicly available from the repositories cited in the manuscript. Processed data are available from the corresponding author on reasonable request.

